# The Incremental Cluster Threshold-Free Cluster Enhancement Algorithm for Functional Connectivity Analysis

**DOI:** 10.64898/2026.04.06.716826

**Authors:** Fabricio Cravo, Raimundo Rodriguez, Alfonso Nieto-Castañón, Stephanie Noble

## Abstract

Threshold-free cluster enhancement (TFCE) is one of the most used statistical inference methods in neuroimaging, but its computational cost limits some of its applications. The current implementations recompute clusters at each threshold step, creating computation costs that poorly scale with precision increases. Furthermore, as larger samples and reduced noise increase maximum t-statistics, computational burden grows correspondingly. As the field moves towards finer parcellations, the number of FC edges grows quadratically with the number of ROIs, making TFCE computationally infeasible at the scales increasingly demanded by the field. We present Incremental Cluster TFCE (IC-TFCE), an algorithm that produces numerically equivalent results to standard TFCE while decoupling runtime from discretization precision. The IC-TFCE builds clusters incrementally from previous threshold steps rather than recomputing them, stores TFCE results on a region of interest (ROI) based structure instead of a functional connectivity (FC) edge structure for improved speed, and can be applied to voxel data through a novel graph transformation described and validated herein. This algorithm achieves a measured 3–93× speedup for FC TFCE depending on the precision parameter *dh*, making TFCE analyses with fine parcellations of 1000 or more ROIs computationally tractable for the first time. Finally, we validate correctness through mathematical proof and numerical comparison. The efficiency provided by IC-TFCE allowed a large-scale empirical power analysis across *dh* values to guide practitioners in parameter selection for their analyses.

## 1 Introduction

Cluster-based inference is one of the most widely used methods for detecting brain activation in functional magnetic resonance imaging (fMRI) [WKW14, Yeu18, LSL^+^18]. Real brain activations are spatially correlated, and cluster methods use this spatial property to improve detection [FWF^+^94, PWEF97, HSY^+^06, WLK07, WKW14]. This clustering is observed in voxel activation patterns [WKW14] and, through their correlations, mathematically implied in functional connectivity data [ZFB12]. The high noise inherent to fMRI [Liu16, BFvG^+^09, VMD^+^21] requires improvements in statistical methods to more reliably detect true effects at realistic sample sizes [SN09, DG02, Bea10].

Two widely used cluster inference methods are cluster size [FWF^+^94] testing and threshold-free cluster enhancement (TFCE) [SN09, SKSN11, SSZ^+^19]. These methods operate on adjacent variables: voxels in activation analyses or edges in FC analyses. For both approaches, a cluster refers to a group of spatially adjacent voxels (for activation data) or connected edges (for FC data) that all exceed a statistical threshold. The cluster size method works as follows: First, a significance threshold is applied to individual variables, forming clusters. Then, a null distribution is estimated by computing the maximum cluster size across permutations of the data. Observed clusters larger than those expected under the null distribution are deemed significant.

The TFCE [SN09] is similar to the cluster size, but it eliminates the need for an arbitrary threshold and has been shown to be a more powerful approach [NSC20, NMZS22]. Instead of testing at a single threshold, TFCE integrates across many thresholds. At each threshold level, it computes the size of the cluster that each variable is contained in. The final TFCE value is a combination of its own statistical strength and the cluster sizes of the clusters this variable is contained in at each threshold level in the form of an integral. In practice, this integral is computed using a discrete approximation to reduce computational burden [JBB^+^12, OFMS11, GLL^+^13, OCH16, WRW^+^14, YCM^+^26]. The approximation uses a step size parameter *dh*. The maximum t-statistic *t*_*max*_ divided by *dh* determines the number of threshold steps *n*_*th*_ = *t*_*max*_*/dh*, with the clusters forming all variables being computed at each threshold step.

However, the TFCE is among the most computationally demanding statistical inference procedures in neuroimaging [SSZ^+^19, FCH^+^22], with documented computational burden limiting its application in some analyses [APE^+^14, SSZ^+^19, FCH^+^22]. Even though some of these limitations address voxel applications, voxel data has at most 26 neighbors per voxel, resulting in a bounded connectivity that greatly reduces computational complexity compared to FC data, where graphs are complete. This becomes particularly relevant for analyses with high *t*_*max*_ values, whether from large sample sizes, high-quality data with reduced noise, or strong effects [BIM^+^13, PBD^+^17, SI20], and for applications requiring finer precision (lower *dh*).

As the field moves towards finer parcellations with greater numbers of ROIs [SKG^+^18, EYG18], the number of FC edges grows quadratically with the number of nodes, rapidly making TFCE computationally infeasible. With 1000 ROIs, for instance, a single FC matrix already contains 499,500 edges. Since TFCE is a permutation-based method requiring on the order of 1000 permutations per analysis [WRW^+^14], the computational burden at this scale becomes prohibitive [SSZ^+^19, FCH^+^22]. If neuroimaging studies continue to move towards even finer parcellations, this problem will worsen, and it will restrict the TFCE application.

Another issue with the current discretized implementation is that it creates a speed-precision trade-off [PLNR15]. Practitioners must choose between using larger *dh* values for faster computation or smaller *dh* values for higher precision. Smaller *dh* requires more threshold steps and therefore more cluster recomputations. While the impact of *dh* [SN09] was already partially analyzed, this analysis was of limited scope and did not assess how it affects statistical power empirically. Some applications could require lower *dh* values than those commonly adopted, but the computational cost makes this impractical. The CONN toolbox [WGNC12] provides an Exact TFCE implementation that eliminates discretization entirely, making *dh* irrelevant. However, this implementation is slower than the discretized TFCE. For sufficiently small *dh*, the precision gains over the discretized TFCE become negligible, yet the Exact implementation cannot take advantage of this speed gain. The field needs an algorithm that provides both speed and precision without forcing a tradeoff.

In this work, we present the Incremental Cluster TFCE (IC-TFCE). The IC-TFCE produces identical results to standard TFCE with a computational speedup that increases as the *dh* becomes smaller. Rather than recomputing all clusters at each threshold step, IC-TFCE builds clusters incrementally from previous iterations. For FC data, we also introduce a node accumulation data structure that enables efficient storage and retrieval of TFCE values for each edge. These strategies reduce computational complexity from *O*(*N*^2^*n*_*th*_) to *O*(*N*^2^ + *n*_*th*_*N*), where *N* is the number of regions of interest (ROIs). Empirically, we observe 3×-93× speedups on FC data depending on the *dh* parameter, type of data, and parcellation used, with greater speedups at smaller *dh* values and with a greater number of nodes. Finally, by reducing computational complexity from 𝒪(*N*^2^*n*_*th*_) to 𝒪(*N*^2^ + *n*_*th*_*N*), IC-TFCE restores the viability of TFCE-based inference at the finer parcellation scales increasingly demanded by the field.

We provide rigorous validation through mathematical proof and numerical comparison with standard TFCE. To guide parameter selection, we conduct a power analysis across different *dh* values, quantifying the precision-power tradeoff that was previously uncharacterized. We provide pseudocode for IC-TFCE implementation, including a graph transformation for voxel data. Finally, we formally describe and validate the Exact TFCE algorithm used in the CONN toolbox [WGNC12], which has not been previously documented in the literature, and provide its pseudocode with incremental clustering strategies.

## 2 Results

### 2.1 The Algorithm

#### 2.1.1 The Original TFCE Algorithm

The TFCE algorithm tests for the statistical significance of variables, either individual voxels or FC edges, based on their ability to form clusters. It does so by transforming each variable’s test statistic (e.g., t-statistics from group comparisons or GLM contrasts) by integrating over cluster sizes at multiple thresholds. Each edge’s TFCE value is computed by integrating the size of clusters it belongs to across threshold levels, giving higher values to edges that form larger connected components.

The TFCE integral requires two parameters: (1) *E*, which weights the extent of clustering, (2) *H*, the height exponent that weights the threshold levels up to each variable’s statistic value. The algorithm’s integral transformation is:

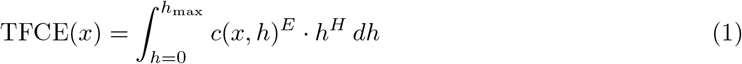

where *c*(*x, h*) is the number of variables (fc edges or voxels) in the cluster containing variable *x* whose test statistic is above threshold *h*.

Normally, this integral is not computed in the previous form, and it is computed with an approximation as a discrete sum:

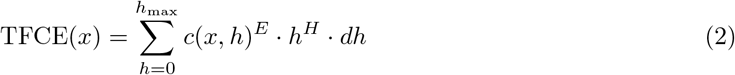

To perform this integration numerically, a new parameter is introduced, *dh*, the step size for discretizing the integral. Like *E* and *H*, the choice of *dh* has historically been empirical, with numerical accuracy of the approximation being the only consideration.

#### 2.1.2 Variables and Topology

The TFCE algorithm is used to two distinct types of neuroimaging data, voxel and functional connectivity, with varying ways of implementation. Furthermore, beyond the data types, the definition of what a cluster (topology) is varies substantially. In this section, we clearly define for which applications the IC-TFCE is oriented towards by explaining the variables and topology used herein.

For voxel activation data, adjacency is defined geometrically: Two voxels are neighbors if they occupy spatially contiguous positions in the brain volume, with varying degrees of neighborhood ranging from 6-connectivity ((side-sharing) to 26-connectivity (side-, diagonal-sharing)) in three dimensions [SN09].

For FC data, the situation is more complex. Each variable is an edge in a connectivity graph and two edges can be declared neighbors in different ways. The Spatial Pairwise Clustering (SPC) [ZFB12] TFCE defines adjacency between edges based on the anatomical proximity of their endpoint ROIs: edges that are connected to spatially adjacent ROIs in the brain are considered neighbors. Another approach, introduced by the Network Based Statistic (NBS) [ZFB10] and later extended to TFCE by Baggio et al. [BAS^+^18], defines two edges as neighbors if they share at least one ROI. Under this node-sharing adjacency, clusters are connected subgraphs of the original ROI graph, and cluster membership for any edge is fully determined by the cluster memberships of its two endpoint nodes.

The IC-TFCE is designed primarily for the NBS-TFCE and can be extended for the voxel TFCE. Therefore, in this manuscript, when we refer to topology for voxels, we refer to their spatial contiguity, and for FC, it signifies that both edges are connected to a common ROI.

#### 2.1.3 The IC-TFCE Algorithm

The IC-TFCE algorithm is simply an alternative way to compute the same TFCE algorithm that yields equal results, but uses the clusters from previous integral thresholds to build the clusters from the following threshold, creating a faster algorithm. The approach processes the clusters from the highest threshold to the lowest, as at the maximum threshold, every variable test statistic is below the threshold, so all clusters are empty. At each step, we add the variables whose test statistics fall between *h*_*n*−1_ = (*n* − 1)*dh* and *h*_*n*_ = *ndh* to compute the next cluster. The incremental merging can be implemented with Union-Find. However, Union-Find provides no complexity gain, as it is required to check the sizes of each cluster for the TFCE computation at each discrete step.

For FC data, to achieve faster complexity, we introduce a node accumulation structure comprising two matrices: (1) **S**, storing cluster sizes as the number of edges at each threshold for each node, and (2) **F**, storing the cumulative TFCE integral values. The structure **S** is populated during cluster merging, then **F** is computed from **S** through a sum. Finally, each edge retrieves its TFCE value in O(1) time by indexing **F** at its threshold bin and either endpoint node. The pseudocode for this implementation is provided in Algorithm 1 with the supporting Figure 1. In Supplementary Section 7, we also prove the following proposition that assures correctness:

##### Proposition 2.1.

*For all discrete integration steps, the IC-TFCE algorithm computes equal clusters to the TFCE*.

For voxel analysis, we provide a graph transformation that converts spatial adjacency into an edge-based representation regardless of dimension. Each edge connects two spatially adjacent voxels and receives the minimum of their test statistics. This allows IC-TFCE to be computed on voxel data, with the only difference being that the TFCE can be calculated directly after each bin is added, without the need for the extra accumulation and edge retrieval steps. The graph transformation is provided in Algorithm 2 and the proof that transformation is indeed correct can be found in the Supplementary Section 8.

For the code implementation of the graph transformation, IC-TFCE for FC, and IC-TFCE voxels, please refer to the PRISME power calculator [CFS^+^26].

#### 2.1.4 The Exact TFCE Implementation

The CONN toolbox [WGNC12] provides an exact TFCE implementation for FC data. Herein, we formalize it and provide a pseudocode implementation of it.

The Exact TFCE works by sorting the variables’ test statistics and varying the *dh* step according to the sorted variables. Since for *h* any between the two test statistics of two sorted variables, the function *c*(*x, h*) remains constant for all variables, one can perform the following integration in piecewise steps.

Let 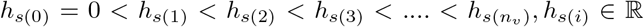 be the sorted test statistics of the variables and *n*_*v*_ ∈ ℕ the number of variables:

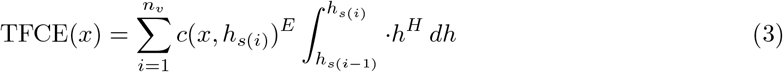

Essentially, the cluster size is constant at those sequential intervals, which allows its removal from the integral and its calculation.

The pseudocode for the Exact TFCE algorithm is presented in Algorithms 4 and 3. Rather than sweeping over all nodes at each edge step, the Exact TFCE maintains a running score per cluster, updated whenever a cluster is accessed. Each cluster tracks its last processed height, so the accumulated TFCE contribution over any interval can be computed in *O*(1) via FlushCluster. When an edge enters, its starting value is recorded as the current cluster score and the final TFCE for that edge is then recovered by walking up the parent chain from its entry cluster and subtracting this starting value. This approach stores one score value per edge rather than an *n*_*th*_ × *N* accumulation matrix, making it more memory efficient than IC-TFCE when *n*_*th*_ > *N*. However, each cluster merge creates a new cluster object, and all node labels in the merged cluster must be updated to point to the new cluster, carrying a higher per-merge cost than the union-by-size strategy used in IC-TFCE. The IC-TFCE can be implemented with a similar strategy.

### 2.2 Computational Complexity

#### 2.2.1 The IC-TFCE Implementation

Note as *N* the number of ROIs. Since a functional connectivity graph is complete, bidirectional and there are no edges from nodes to themselves, the number of edges *E* is 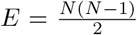. Let *n*_*th*_ be the number of threshold bins which is *dh* divided by the maximum t-stat in the graph.

From the pseudocode, the IC-TFCE complexity is the following:

##### Algorithm 1 IC-TFCE for FC Data

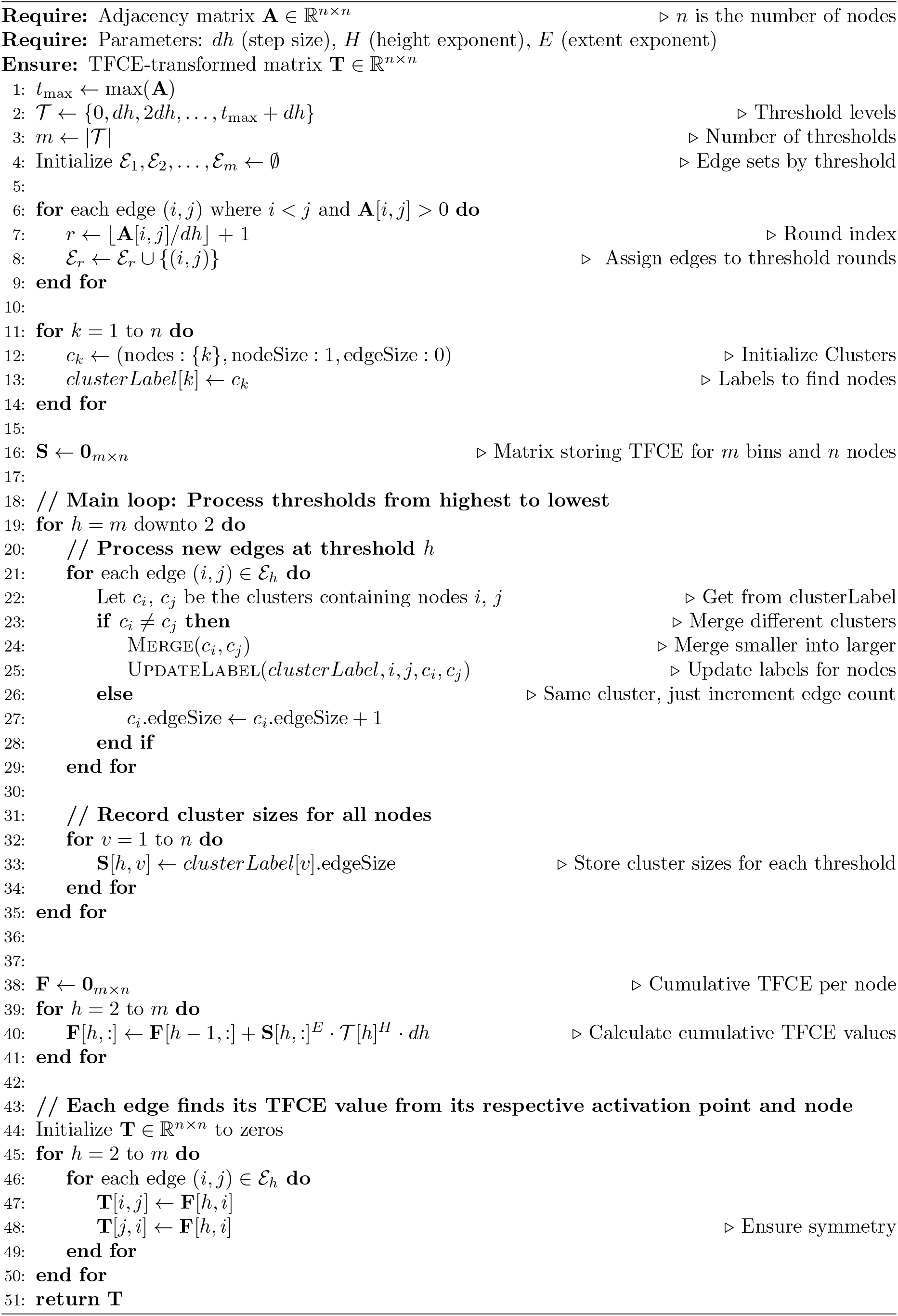

**Figure 1:**
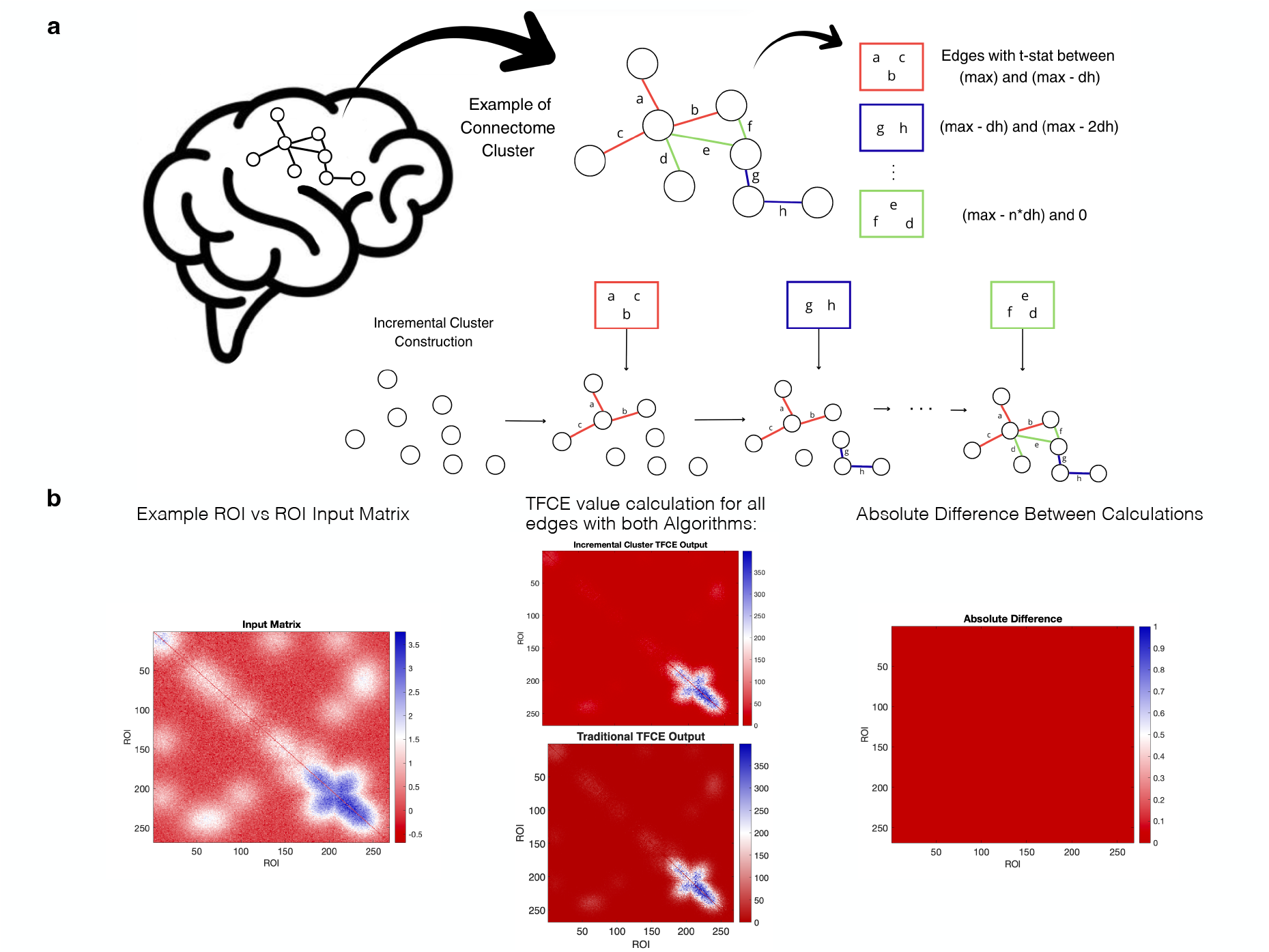
TFCE algorithm comparison and validation. **(a)** Figure describing the IC-TFCE Algorithm through a visual application using an example connectome cluster from functional connectivity data. Edges are categorized by t-statistic threshold ranges (red: maximum to max-dh; blue: max-dh to max-2dh; green: max-n*dh to 0). The edges then form a cluster incrementally for the TFCE computation. **(b)** Numerical comparison between TFCE implementations. Left: Example ROI-to-ROI input matrix from resting-state functional connectivity. Middle: TFCE values computed using the IC-TFCE and the TFCE show nearly identical results. Right: Absolute difference between calculations demonstrates numerical equivalence (difference *<* 0.001 for all edges).

- Assigning each edge to their respective bin 𝒪(*N*^2^)
- The IC-TFCE algorithm processes each edge once and it is of *O*(*N*^2^) order. For each edge, either: the edge connects nodes already in the same cluster and is processed in *O*(1) time, or it connects nodes in different clusters, triggering a merge. Since each merge reduces the cluster count by one and we begin with *N* singleton clusters, at most *N* − 1 merges occur across the entire algorithm. Since we merge smaller clusters into larger ones, in the best case the largest cluster absorbs a single node at each step for a total merge cost of *O*(*N*). Differently than the Union-Find algorithm [Tar75, TvL84, CF07], we update cluster labels for the entire cluster at each merger as they are necessary for the cluster size calculation for each threshold. On the worst case merge, clusters start as single nodes and double in size every time they merge. A level number *l* indicates the number of the stage where clusters have size 2^*l*^. At level *l*, there are *N/*2^*l*^ clusters of size 2^*l*^, which go through *N/*2^*l*+1^ merges where each merge costs *O*(2^*l*^). The cost at each level is therefore 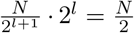. Since there are log_2_ *N* levels, the total merge cost is:

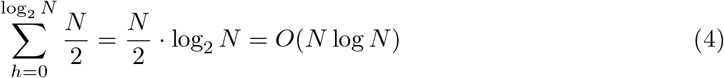 #### Algorithm 2 Graph conversion

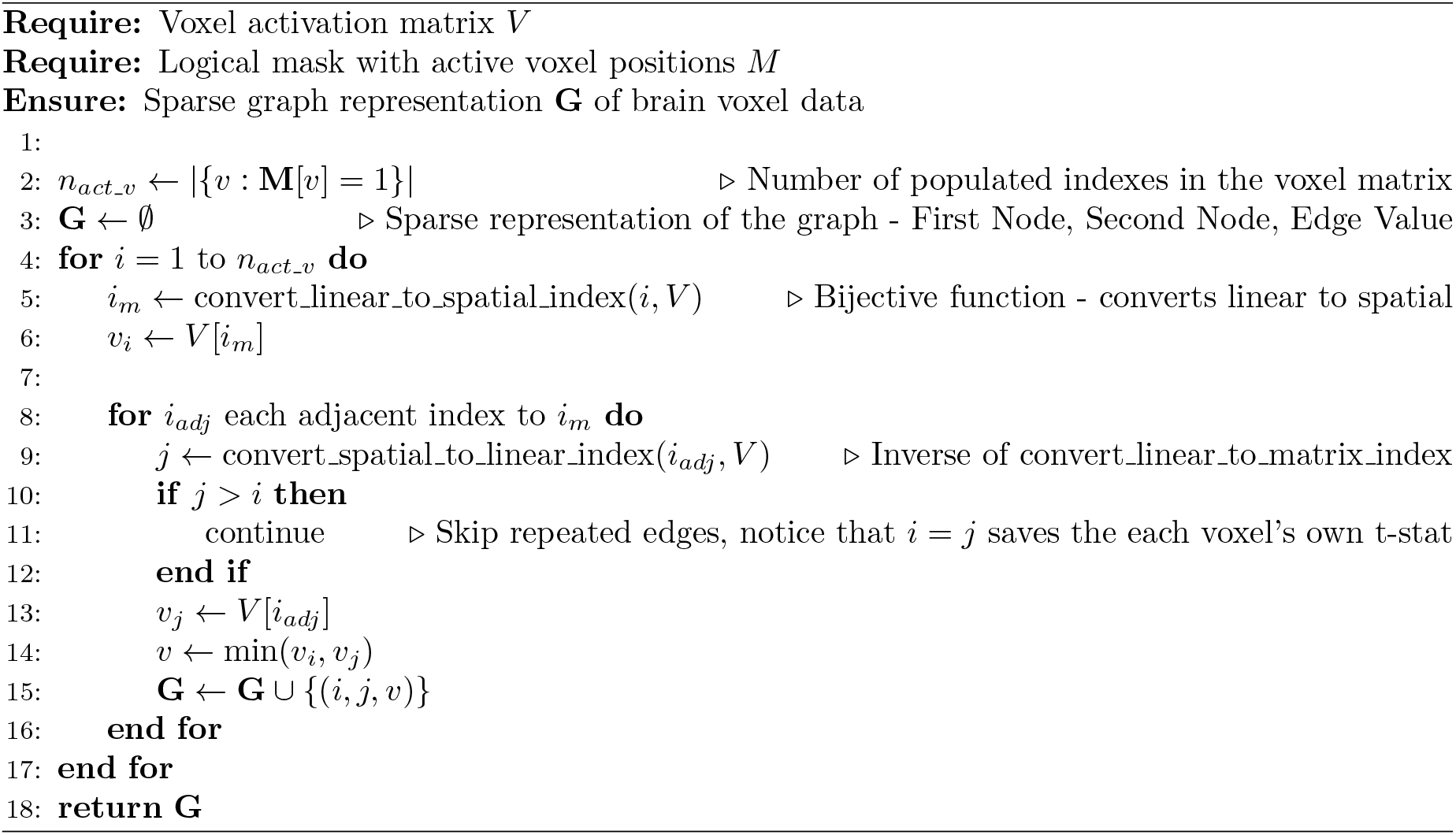 Since we process each edge once, the total cost over all edges is:

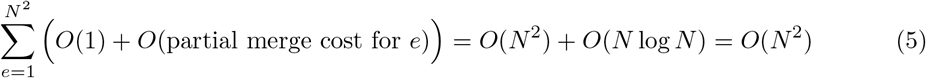

where the partial cost for merging is zero when an edge *e* simply gets added to the cluster with no merger. At the end of each threshold step, we record the cluster size for each of the *N* nodes, for a total cost of *O*(*n*_*th*_ · *N*). Therefore, the cost of the

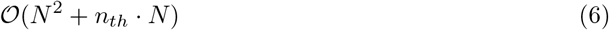
- Accumulating the TFCE value for each node for each threshold bin is performed in 𝒪(*n*_*th*_*N*) time.
- Finding the TFCE value for each edge for each edge is performed with a single check per edge by checking the structure that accumulated the cluster size in 𝒪(*N*^2^).

Therefore, by summing through all complexities, the overall complexity of the IC-TFCE is 𝒪(*N*^2^ + *n*_*th*_*N*). On the other hand, the traditional TFCE algorithm computational complexity is 𝒪(*N*^2^*n*_*th*_).

#### 2.2.2 The IC-TFCE Exact Implementation

The IC-TFCE accumulation strategy can also be applied to compute the Exact TFCE. Rather than using the discretized thresholds, the edges are sorted by their test statistics and each edge defines a single threshold bin. The number of threshold bins is therefore equal to the number of edges, 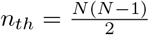, which is of *O*(*N*^2^) order. Since each edge’s TFCE value depends only on the cumulative integral at its two endpoint nodes, the accumulation structure can be replaced by storing the initial TFCE value at the time each edge enters and subtracting it from the final cumulative value at the corresponding node. The overall complexity then follows directly from the IC-TFCE formula:

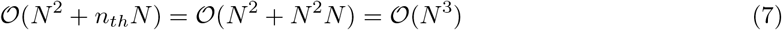

##### Algorithm 3 Exact TFCE Helper Functions

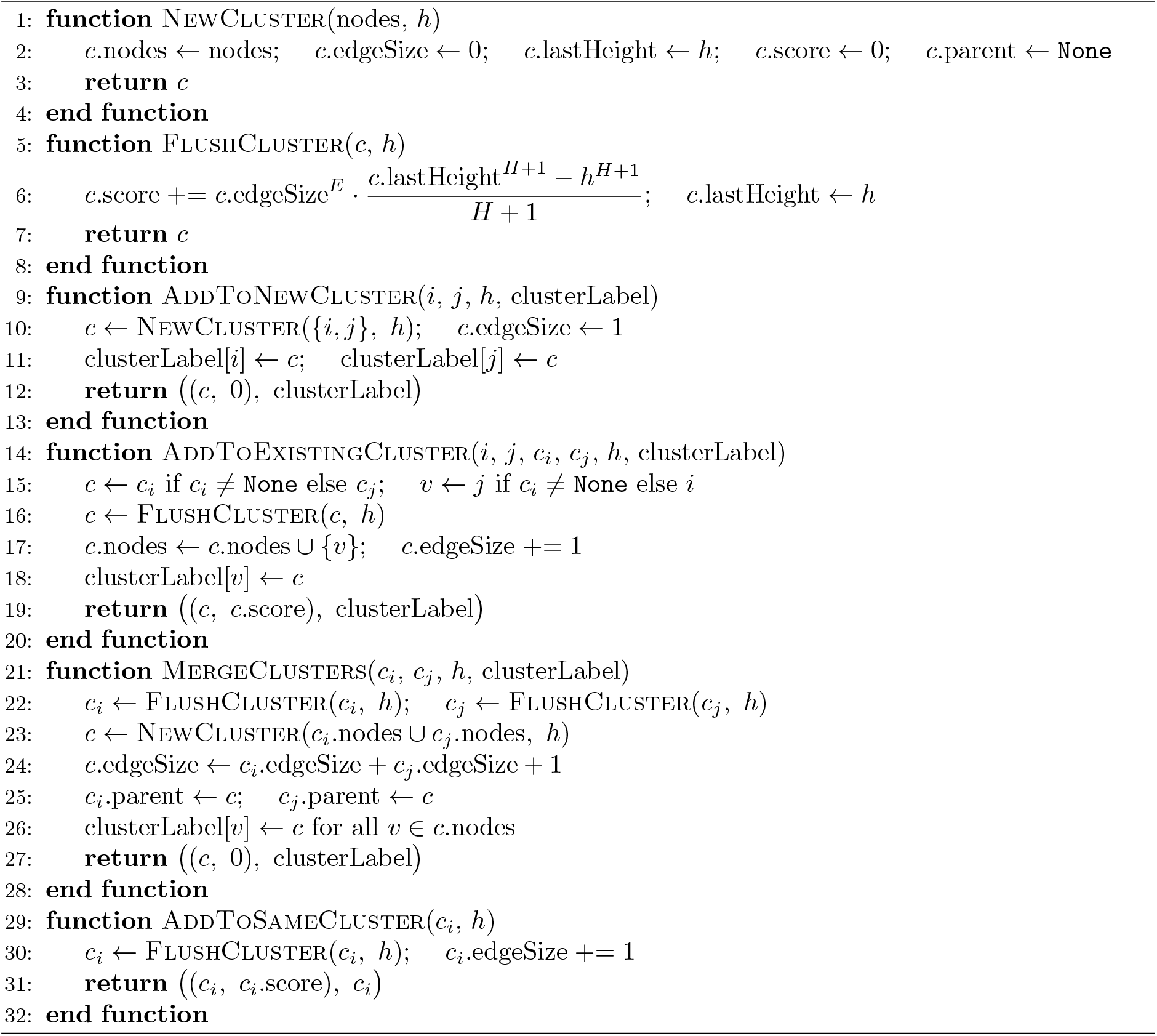

##### Algorithm 4 Exact TFCE

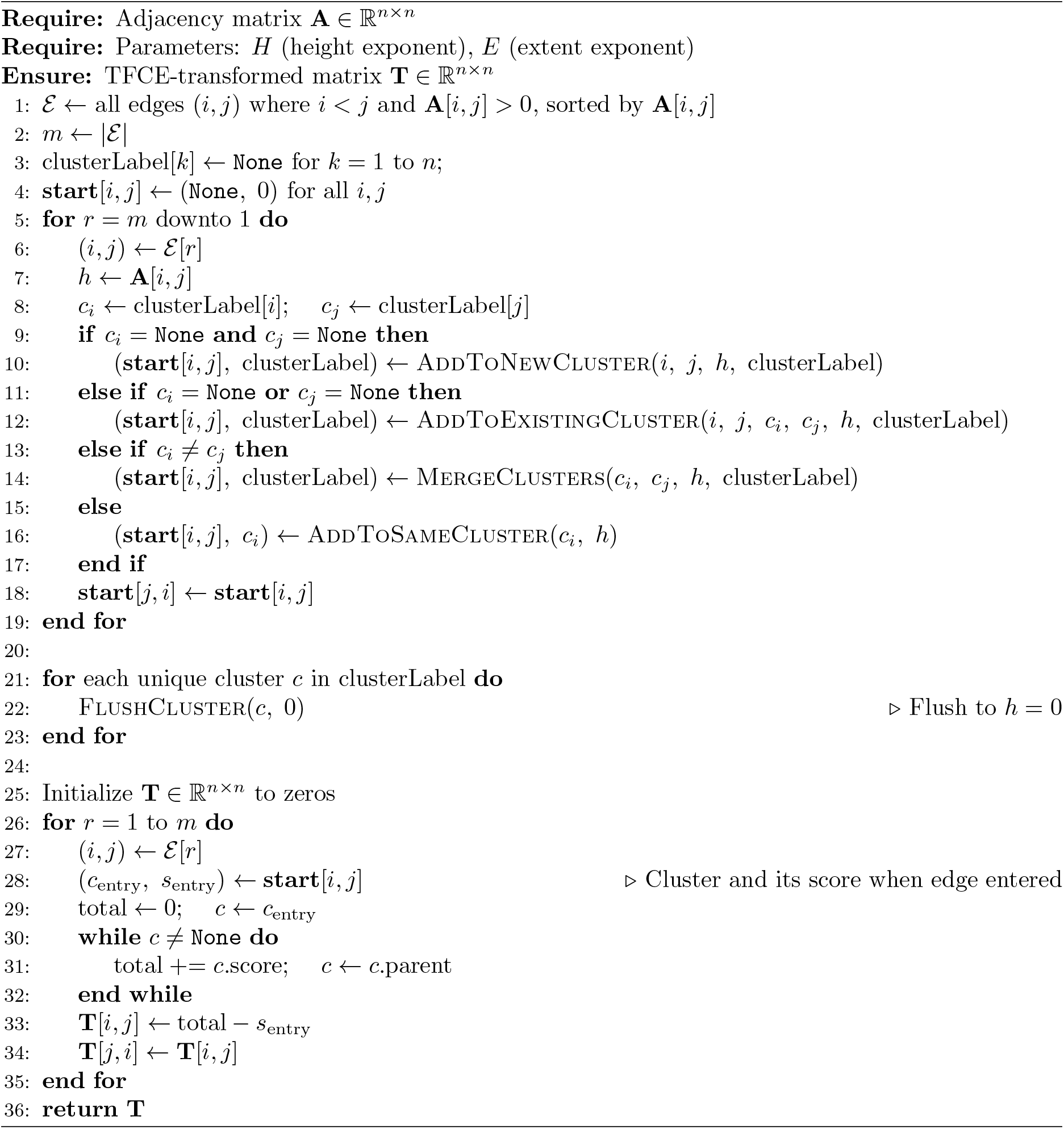

#### 2.2.3 Voxel Transformation Complexity

The graph transformation from voxel data requires iterating over all voxels and their adjacency relationships, yielding a complexity of:

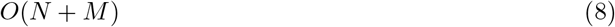

where *N* is the number of voxels and *M* is the number of adjacency links between neighboring voxels. In voxel data, each voxel has a bounded degree with at most 26 neighbors under full 3D adjacency. Therefore, *M* = *O*(*N*), and the transformation complexity becomes:

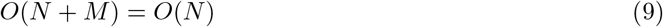

For voxel data, the accumulation structure is no longer necessary. Cluster sizes are defined by the nodes themselves and the TFCE values are updated after each merge. Following analogous analysis to the FC case, the IC-TFCE complexity for voxel data comprises:

- Edge bin assignment: *O*(*M*) = *O*(*N*)
- Processing edges and merges: *O*(*M* + *N* log *N*) = *O*(*N* log *N*)
- Recording cluster sizes at each threshold step: *O*(*n*_*th*_ · *N*)

The overall complexity of IC-TFCE for voxel-based analysis is therefore:

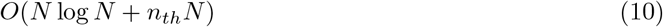

Due to the bounded number of edges, traditional TFCE on voxel data has complexity *O*(*N* ·*n*_*th*_), which is asymptotically equivalent as log_2_ *N* ≤ *n*_*th*_ for the majority of cases. For example, with *N* = 500 000 voxels, log_2_ *N* ≈ 19, while even with coarse precision (*t*_*max*_ = 10, *dh* = 0.25), *n*_*th*_ = 40. The graph transformation therefore provides a unified implementation framework for both FC and voxel analyses using the IC-TFCE. However, the IC-TFCE benefit is clearly stronger in FC-based analysis.

From the mathematical proofs in Supplementary Section 8, the transformation still remains valuable to allow for graph based algorithms to be applied directly to voxel data, as clusters in this graph have exactly the same voxels as the clusters as defined from voxel data. Furthermore, this allows the Exact TFCE to be used in voxel analysis.

### 2.3 Numerical Simulations

Even though the IC-TFCE has better computational complexity than the traditional TFCE implementation, it does not necessarily imply the algorithm is faster in practical applications. The IC-TFCE has more overhead costs related to preparing data structures in the form of the edge bin assignment and preparing the node accumulation structure.

To guarantee the IC-TFCE provides a substantial speed gain, specially for cases where *dh* is smaller, we measure the computational gain in speed under the same populational conditions. Since the t-statistics of the edges themselves affects the execution speed of the TFCE algorithm, we perform an analysis for multiple subject group numbers (20, 80, 200) over five different tasks of the HCP dataset: working memory, motor, language, social, and relational [CFS^+^26]. As the idea is to focus on speed difference instead of inference, only 50 permutations are used and 10 repetitions for each task for each subject group number.

As we can see (Table 1), the IC-TFCE implementation results has the lowest mean time execution across all tasks on the HCP dataset and this result can be observed over multiple subject sizes and over the range of currently recommended *dh* values used in literature. Even for the highest *dh* of 0.25, we still observe a significant 3× times speedup. The Exact TFCE, although it yields the most precise computation of all TFCE algorithms, is by far the slowest one.

**Table 1:** Average runtime comparison of TFCE implementations across varying sample sizes for a group-level analysis. All algorithms were implemented in C++ to ensure fair performance comparison. Values represent mean execution times per permutation, averaged across five tasks and 10 repetitions from the HCP dataset (500 total executions per condition).

| **(a) Subject Numbers = 20** |  |  |  |
| --- | --- | --- | --- |
| $dh$ | IC-TFCE (ms) | TFCE (ms) | Speedup |
| 0.01 | 4.024 | 133.186 | 33.10x |
| 0.05 | 2.347 | 30.037 | 12.80x |
| 0.1 | 2.243 | 16.424 | 7.32x |
| 0.25 | 2.238 | 7.228 | 3.23x |
| **(b) Subject Numbers = 80** |  |  |  |
| $dh$ | IC-TFCE (ms) | TFCE (ms) | Speedup |
| 0.01 | 3.890 | 121.781 | 31.30x |
| 0.05 | 2.395 | 27.794 | 11.61x |
| 0.1 | 2.115 | 14.607 | 6.90x |
| 0.25 | 2.013 | 6.587 | 3.27x |
| **(c) Subject Numbers = 200** |  |  |  |
| $dh$ | IC-TFCE (ms) | TFCE (ms) | Speedup |
| 0.01 | 3.946 | 121.847 | 30.88x |
| 0.05 | 2.522 | 27.601 | 10.94x |
| 0.1 | 2.264 | 14.578 | 6.44x |
| 0.25 | 2.027 | 6.606 | 3.26x |

To evaluate how computational gains scale with the number of ROIs, we conducted an additional benchmark using synthetic data. The ROI-by-ROI t-statistic matrices were generated for varying numbers of ROIs by sampling t-statistics from a standard normal distribution and applying a 2D Gaussian filter to introduce spatial correlation structure expected in fMRI. Table 2 presents the runtime comparison between IC-TFCE and traditional TFCE across ROI counts ranging from 200 to 1000, demonstrating that IC-TFCE’s computational advantage grows with the number of ROIs.

**Table 2:** Average runtime comparison of TFCE implementations across varying numbers of ROIs on synthetic data. Synthetic t-statistic matrices were generated by sampling from a standard normal distribution, applying a 2D Gaussian filter (zero-padded, *σ* scaled proportionally to *N*) to introduce spatial correlation structure, and symmetrizing the result. Times represent mean execution across 500 repetitions. Spatial correlation structure size was kept constant across different parcellations. All algorithms were implemented in C++.

| (a) N = 200 ROIs |  |  |  |
| --- | --- | --- | --- |
| $dh$ | IC-TFCE (ms) | TFCE (ms) | Speedup |
| 0.01 | 0.9 | 33.2 | 35.3x |
| 0.05 | 0.3 | 6.6 | 20.9x |
| 0.1 | 0.2 | 3.3 | 14.4x |
| 0.25 | 0.2 | 1.4 | 7.6x |
| (b) N = 500 ROIs |  |  |  |
| $dh$ | IC-TFCE (ms) | TFCE (ms) | Speedup |
| 0.01 | 2.8 | 183.7 | 64.5x |
| 0.05 | 1.3 | 36.9 | 28.9x |
| 0.1 | 1.1 | 18.7 | 17.4x |
| 0.25 | 0.9 | 7.5 | 7.9x |
| (c) N = 1000 ROIs |  |  |  |
| $dh$ | IC-TFCE (ms) | TFCE (ms) | Speedup |
| 0.01 | 7.5 | 703.6 | 93.9x |
| 0.05 | 4.1 | 135.6 | 33.0x |
| 0.1 | 3.6 | 68.3 | 18.7x |
| 0.25 | 3.4 | 28.7 | 8.3x |

Table 2 shows that IC-TFCE benefits scale with parcellation sizes. Speedups increase substantially with the number of nodes, reaching up to 93.9× at *N* = 1000 ROIs with *dh* = 0.01, compared to 7.6× at *N* = 200 with *dh* = 0.25. As expected from the complexity analysis, finer *dh* values yield the greatest relative gains, since the traditional TFCE runtime grows proportionally with *n*_*th*_ while IC-TFCE does not. Even the most conservative case (*dh* = 0.25) delivers consistent speedups over all analysis. Given that larger parcellations and finer precision are precisely the conditions under which TFCE has historically been computationally slower, IC-TFCE makes previously infeasible analyses tractable without any sacrifice in numerical accuracy.

### 2.4 Power Analysis

With our algorithmic optimizations, we conducted a large scale empirical power analysis using data from the Human Connectome Project [VESB^+^13]. For functional connectivity analysis, we tested five task contrasts across varying sample sizes (*n*_*sub*_ = 20, 40, 80, 120, 200) and four *dh* values (0.01, 0.05, 0.1, 0.25). Statistical power was calculated using the PRISME [CFS^+^26] power calculator. We examined two power metrics: (1) overall average power, calculated as the mean power across all edges in the connectivity matrix, and (2) network-based average power, calculated as the mean power across-task of the top ranked networks in in-task power. For voxel activation analysis, we performed similar comparisons using the same *dh* parameters and 3 sample sizes (*n*_*sub*_ = 20, 80, 200). Results demonstrated that statistical power increased with sample size for all *dh* values, but differences between *dh* parameters were non-existent, particularly at larger sample sizes. Both overall average power and network-based average power showed consistent patterns across different *dh* values at *n*_*sub*_ = 80 and *n*_*sub*_ = 200, with only modest variations observed at the smallest sample size (*n*_*sub*_ = 20).

## 3 Methods and Data Processing

The power analysis was performed using the PRISME empirical power calculator [CFS^+^26] with *E* = 0.5 and *H* = 2 for voxel data and *E* = 0.4 and *H* = 3.0 for FC data. Power analysis used 800 permutations for the TFCE calculations and 100 repetitions for calculating power. The 800 permutation usage is close enough in precision to the 1000 normally used and it is used to speed up the large scale analysis. The 100 repetitions usage provides approximately ±5% precision on power estimates, sufficient for practical sample size planning. From PRISME algorithm, all TFCE implementations were compared across identical permutation and subject data for each repetition, relative power differences between dh values are estimated with higher precision.

Functional images were motion-corrected using SPM5. The data were then iteratively smoothed to an equivalent smoothness of a 2.5 mm Gaussian kernel in order to ensure uniform smoothness across the dataset. White matter and CSF were defined on a MNI-space template brain and eroded in order to minimize inclusion of grey matter in the mask. The template was then warped to subject space using a series of transformations described in the next section. This ensured that mainly grey matter voxels were used in subsequent analyses. The following noise covariates were regressed from the data: linear, quadratic, and cubic drift, a 24-parameter model of motion, mean cerebrospinal fluid signal, mean white matter signal, and mean global signal. Finally, data were temporally smoothed with a zero mean unit variance Gaussian filter (cutoff freq=0.19 Hz). Anatomical data were first skull-stripped using FSL. Functional data for each subject, scanner, and session were linearly registered to the corresponding FLASH images. FLASH images were then linearly registered to MPRAGE images. Next, an average MPRAGE image for each subject was created by linearly registering and averaging all 4 anatomical images (from 2 scanner × 2 sessions) for each subject. These average MPRAGE images were used for an iterative nonlinear registration to MNI space. The use of the average anatomical images and a single nonlinear registration for each subject ensures that any potential anatomical distortions due to the different scanners does not introduce a systematic bias into the registration. The average MPRAGE images were nonlinearly registered to an evolving group average template in MNI space as described previously. All transformation pairs were calculated independently and then combined into a single transform that warps single participant results into common space. From this, all subjects’ images can be transformed into common space using a single transformation, which reduces interpolation error. [NST^+^17]

**Figure 2:**
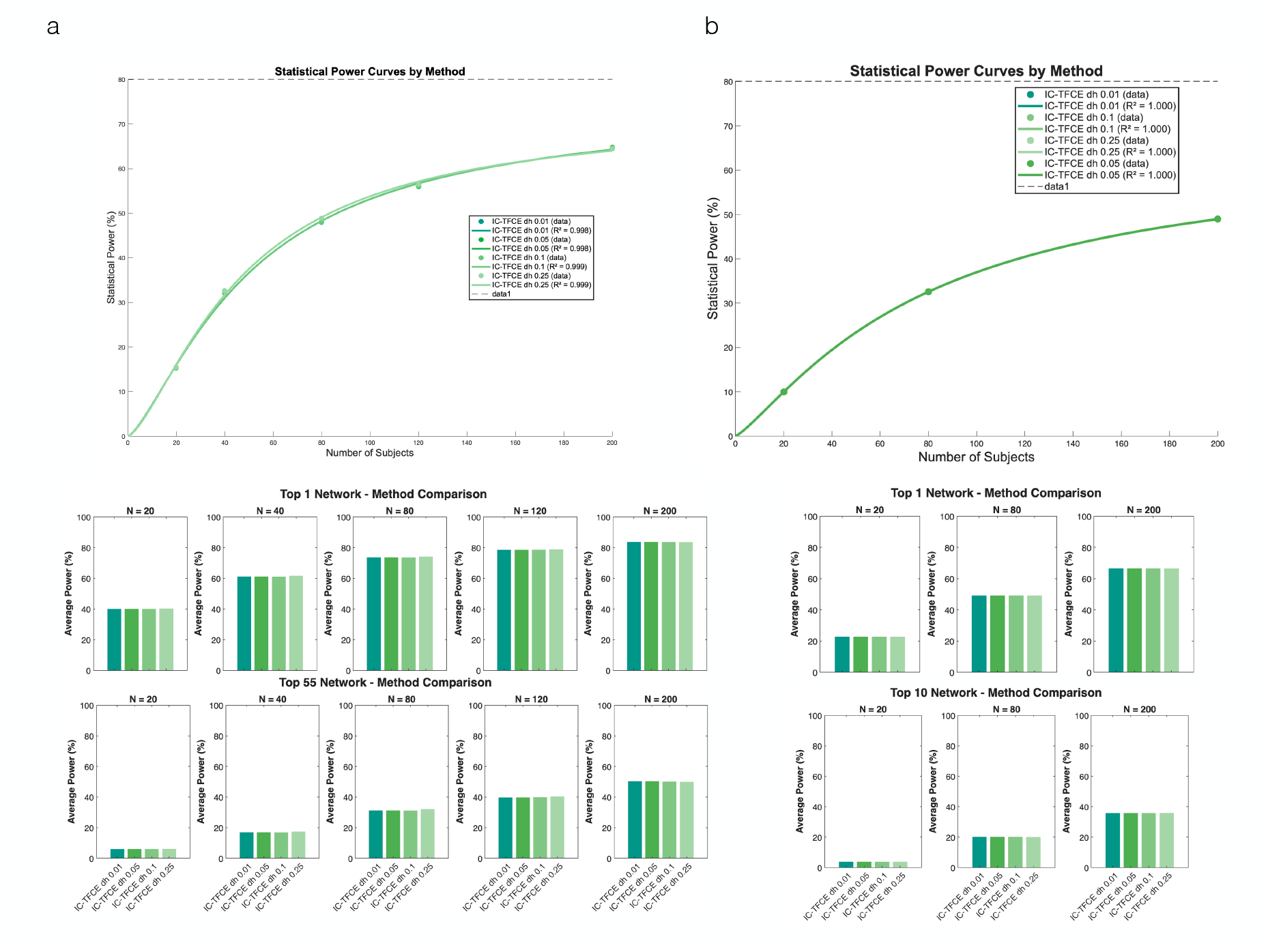
Statistical power comparison of IC-TFCE implementations across different step sizes (dh).: (a) Functional connectivity power analysis showing statistical power curves (top) and average power for top networks (middle and bottom) across varying sample sizes (N = 20, 40, 80, 120, 200) and dh values (0.01, 0.05, 0.1, 0.25). (b) Voxel-based power analysis demonstrating similar trends for IC-TFCE with different dh parameters (0.01, 0.05, 0.1, 0.25). Both analyses show that power increases with sample size, with minimal differences between dh values, particularly at larger sample sizes. Bar charts display average power for top-ranked networks/clusters at N = 20, 80, and 200.

## 4 Discussion

We present IC-TFCE, an incremental cluster algorithm that produces results equal to those of the TFCE, but with improved computational efficiency. The algorithm builds clusters incrementally across threshold levels by reusing clusters from previous steps instead of recomputing them at each threshold. For FC, the traditional TFCE has computational complexity *O*(*N*^2^*n*_*th*_), where N is the number of ROIs or voxels, and *n*_*th*_ is the number of threshold bins (the maximum t statistic divided by the TFCE integration step size *t*_*max*_*/dh*). IC-TFCE reduces this to *O*(*N*^2^ + *n*_*th*_*N*), making the runtime less dependent on discretization precision. On FC data, this translates to measured speedups of 3–93× across multiple dh values, with greater advantages at finer precision. The method extends to voxel activation data through a graph-based transformation that unifies both data types.

Due to the increased computational efficiency of the IC-TFCE for multiple values of *dh*, we can now perform empirical power analysis on different values of *dh*, which were computationally impractical with the traditional TFCE. In this power analysis, we compared the overall average power and the average power of the top-most powered network and found that increasing the precision (decreasing *dh*) does not provide meaningful increases in average power and per network average power. Our power analysis found no meaningful benefit to increasing precision beyond *dh* = 0.1, consistent with current recommendations [SN08]. For large-scale analyses, the value *dh* = 0.25 could be justified. Furthermore, even in analysis using *dh* = 0.25, the speed up provided by the IC-TFCE (3×) is still substantial compared to the traditional implementation.

While our power analysis supports *dh* = 0.25, we focus on recommendation to large-scale applications. First, because this evidence was generated on specific HCP task contrasts and may not generalise to all study types. Second, unless computational constraints make finer precision impossible or impractical, we recommend maintaining *dh* = 0.1 or lower, as there is no reason to sacrifice precision unnecessarily. For novel applications where the appropriate *dh* is uncertain, practitioners who wish to justify increased values of *dh* could compare results with the Exact TFCE for validation on a small subset of data to assess if the added error provides a meaningful difference.

For group-level analysis, since higher *n*_*sub*_ increases the observed *t*_*max*_, which in turn increases *n*_*th*_ with fixed *dh*, one could expect IC-TFCE’s advantage over traditional TFCE to grow with sample size. Table 1 shows that in practice this effect is minimal: execution times remain nearly flat across *n*_*sub*_ = 20, 80, 200 for both algorithms, and the speedup ratios are correspondingly stable. This can be explained by the permutations, as the computation time of the TFCE is usually governed by the 1000 permutations, as opposed to the single data computation. The permutation null distribution, rather than the observed statistics, has a lower effective *t*_*max*_ than the significant data, and the increase of *t*_*max*_ for the permutation data is not as great as the increase of the data itself, as evidenced by the increased rejections with increased sample sizes. The label shuffling from the permutation distribution redistributes the signal across subjects, preventing the permutation *t*_*max*_ from growing substantially with the sample size. Practitioners can therefore expect consistent speedups from IC-TFCE regardless of cohort size.

We can estimate the practical impact of IC-TFCE with two examples. For an individual analysis with *N* = 1000 ROIs and 10,000 permutations, traditional TFCE requires 10000 × 68.3 ms ≈ 11 min per run and the IC-TFCE reduces this to 10000 × 3.6 ms ≈ 36 s. In practice, analyses must be rerun multiple times to verify preprocessing steps, check parameter choices, and correct errors, turning what would be a half-day of iteration into minutes. On larger systematic sensitivity analyses, this implementation is vital. Baggio et al. [BAS^+^18] tested 260 combinations of *E* and *H* parameters for FC-TFCE; completing such a search at *N* = 1000 ROIs with 10,000 permutations would require 260 × 11 min ≈ 49 hours under traditional TFCE, but only 260 × 36 s ≈ 2.6 hours under IC-TFCE.

Although IC-TFCE improves sequential code execution, traditional TFCE offers potential parallelization advantages in specific scenarios. Traditional TFCE can parallelize cluster computation across thresholds as each is computed independently. However, as TFCE is used as a permutation test requiring thousands of executions, parallelizing across permutations is the current standard. The IC-TFCE supports the permutation-level parallelization strategy. Furthermore, parallelizing both within-execution cluster computation and across permutations would require computational resources beyond what is typically available. The IC-TFCE’s faster single-execution time still provides practical advantages even when parallelization is available.

The IC-TFCE addresses a scaling challenge and a computational resource challenge. It is already outperforming commonly used TFCE implementations and is necessary for conducting large-scale studies. By reducing computational complexity from 𝒪(*N*^2^*n*_*th*_) to 𝒪(*N*^2^ + *n*_*th*_*N*) for FC analysis, IC-TFCE delivers better computational efficiency while maintaining numerical equivalence to traditional implementations. Our power analysis validates that the standard precision used for integration (*dh* = 0.1) provides adequate statistical power. As neuroimaging continues toward larger sample sizes and better signal quality, IC-TFCE’s computational advantages are predicted by statistical theory to increase, making it the appropriate implementation for TFCE-based inference in both functional connectivity and voxel activation analyses. An implementation of the algorithm is available in the PRISME toolbox, and the code can be adapted to existing toolboxes.

## 5 Data and Code Availability

The IC-TFCE implementation and all analysis code are available in the PRISME toolbox [CFS^+^26] under the TFCE directory (https://github.com/neuroprismlab/PRISME-Brain-Power-Calculator). Data used in this study are from the Human Connectome Project [VESB^+^13], available at https://www.humanconnectome.org subject to the HCP

## 6 Acknowledgments

This work was supported by the National Institute of Mental Health (K99 MH130894 and R00 MH130894 to Stephanie Noble). Data were provided by the Human Connectome Project, WU-Minn Consortium (Principal Investigators: David Van Essen and Kamil Ugurbil; 1U54MH091657) funded by the 16 NIH Institutes and Centers that support the NIH Blueprint for Neuroscience Research; and by the McDonnell Center for Systems Neuroscience at Washington University.

## 7 TFCE and IC-TFCE Equality

Let *G* = (*N, E, W*) be an undirected weighted graph generated from the calculated *t*-statistics from a GLM fit, with *N* being the set of nodes, *E* being the set of edges, and *W* a function from *E* → ℝ that returns a *t*-statistic associated with an edge *e* ∈ *E*.

We assume the existence of at least one positively weighted edge in *G*:

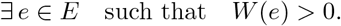

and that the number of edges is finite:

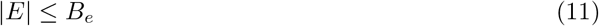

with *B*_*e*_ ∈ ℕ_>0_ and |*E*| the number of edges.

Furthermore, we assume the function *W* is bounded, so it maps all finite number of edges to finite values. Therefore, there exists a *B*_*w*_ ∈ ℝ such that for all *e* ∈ *E*:

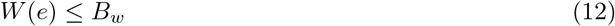

As a final assumption, we assume the graph is undirected: if (*n*_*i*_, *n*_*j*_) ∈ *E*, then (*n*_*j*_, *n*_*i*_) ∈ *E* and *W*(*n*_*i*_, *n*_*j*_) = *W*(*n*_*j*_, *n*_*i*_). Consequently, paths in *G* are reversible. If there exists a path from node *n*_*a*_ to node *n*_*b*_ implies the reverse path from *n*_*b*_ to *n*_*a*_ is also in the graph. The assumption holds for the symmetric functional connectivity graphs. For simplification, this assumption will remain implicit for the extent of this proof and paths will be considered symmetric.

Let δ*t* > 0, δ*t* ∈ ℝ be the integration step size of the TFCE algorithm.

The TFCE algorithm computes the following: the subgraphs given a cut off threshold and the clusters from that subgraph. Finally, with the cluster sizes, the TFCE integral is calculated. We start by defining the subgraphs calculated by the TFCE and then it’s clusters.

### TFCE subgraphs

Let *k* ∈ ℕ. The sequence of subgraphs (*G*_*k*_)_*k*∈ℕ_ with *G*_*k*_ = (*N, E*_*k*_, *W*) where *E*_*k*_ is a subset of edges such that

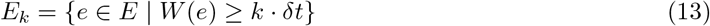

is the sequence of subgraphs calculated by the TFCE algorithm.

### TFCE clusters

The clusters for the TFCE are the maximum connected induced subgraphs (MCISs) of the subgraph *G*_*k*_ = (*N, E*_*k*_, *W*). We now define a maximum connected induced subgraph.

If 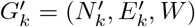 is an MCIS from *G*_*k*_ = (*N, E*_*k*_, *W*), it is:

- For any two nodes 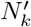, there exists a path between them using only edges in 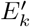,
- No additional node from 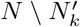 can be added to 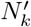 without violating this property (i.e. the number of nodes is maximal).
- 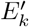 contains all edges from *E*_*k*_ that connect pairs of nodes in 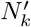 (i.e., *E*^*′*^ = {*e*_*i,j*_ ∈ *E*^*′*^ | *n*_*i*_, *n*_*j*_ ∈ *N*^*′*^}) (i.e. the number of edges is maximal).

To prove the IC-TFCE produces equal results, we need to define both the subgraphs and clusters generated by the IC-TFCE and then show that they are equal to the ones produced by the TFCE. We will start by defining the subgraphs generated by the IC-TFCE and show they are equal. Finally, we will do the same for the clusters.

### IC-TFCE Subgraphs

Now, we define the subgraphs generated by the IC-TFCE implementation.

We start with some set definitions.

Let 𝒯 = (*T*_*k*_)_*k*∈ℕ_ and ℋ = (*H*_*k*_)_*k*∈ℕ_ be sequences of sets indexed by *k* ∈ ℕ.

Let each *T*_*k*_ ∈ 𝒯 be the set of edges defined as follows:

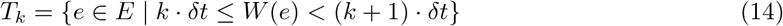

Further, let *H*_*k*_ be the sets defined at such:

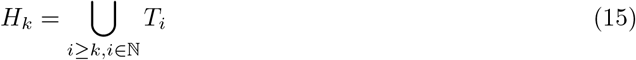

The sequence of subgraphs generated by the IC-TFCE algorithm (*I*_*k*_)_*k*∈ℕ_ are the subgraphs with *I*_*k*_ = (*N, H*_*k*_, *W*).

We first prove a Lemma that will be used by most induction steps in this proof. It is used to establish the base cases.

#### Lemma 7.1.

*There exists a K* ∈ ℕ *such that E*_*k*_ = *H*_*k*_ = ∅ *for k* ≥ *K*

*Proof*. Let *w*_*max*_ = *max*_*e*∈*E*_*W*(*e*). The maximum is well defined because the graphs are finite and the function *W* is bounded. Let *K* ∈ ℕ be the lowest number such that *K* · δ*t* > *w*_*max*_.

From the definition of *E*_*k*_, one has that

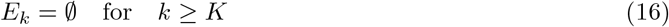

Furthermore, for all *e*, it is *W*(*e*) *< K*δ*t*. Therefore, from the definition of *T*_*k*_, one has that *T*_*k*_ = ∅ for *k* ≥ *K*. It follows from the definition of *H*_*k*_ that *H*_*k*_ = ∅

It is *E*_*k*_ = *H*_*k*_ = ∅ for *k* ≥ *K* and the proof is concluded. □

#### Lemma 7.2.

*For all k* ∈ ℕ, *we have E*_*k*_ = *H*_*k*_.

*Proof*. To prove this, one can show that assuming an element belongs to *E*_*k*_ implies it must be in *H*_*k*_, and vice-versa (i.e. mutual inclusion).

- **Let** *e* ∈ *H*_*k*_: Since *e* ∈ *H*_*k*_, there exists a *T*_*i*_ with *i* ≥ *k* such that *W*(*e*) ∈ [*i* · δ*t*, (*i* + 1) · δ*t*), therefore, we can conclude *W*(*e*) ≥ *i* · δ*t* and consequently *e* ∈ *E*_*k*_.
- **Let** *e* ∈ *E*_*k*_: Since *e* ∈ *E*_*k*_, one has that *W*(*e*) ≥ *k* · δ*t*. From Lemma 7.1, one can conclude *W*(*e*) ∈ [*k* · δ*t, K*δ*t*). Therefore, since the sets {[*i* · δ*t*, (*i* + 1) · δ*t*)}_{*i*∈ℕ|*i*≥*k,i*≤*K*}_ form a disjoint partition of [*k* · δ*t, K*δ*t*), there exists an index *i* ∈ ℕ such that *W*(*e*) ∈ [*i* · δ*t*, (*i* + 1) · δ*t*), and *e* ∈ *T*_*i*_ with *i* ≥ *k*. Since *e* ∈ *T*_*i*_, *i* ≥ *k*, one has that *e* ∈ *H*_*k*_.

It follows by subset inclusion that *H*_*k*_ = *E*_*k*_.

#### Corollary 7.3.

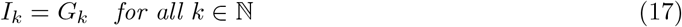

*Proof*. This is a direct consequence of Lemma 7.2 that establishes an equality between edges. From the definition of *I*_*k*_, it has the same nodes and the same weight function for edges as *G*_*k*_. Therefore, the subgraphs are equal. □

Now all that is left to prove is that the IC-TFCE truly computes the correct clusters at each iteration. We start by defining the clusters outputted by the algorithm and then show they are indeed maximum connected induced subgraphs.

#### Definition 7.4.

(IC-TFCE Cluster Construction). Let *C*_*k*_ be the set of clusters produced by the ICTFCE from the subgraph *G*_*k*_, indexed by nodes *x* ∈ *N* :

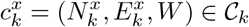

constructed by backwards induction from *K* to 0 as follows:

- **Base case**: 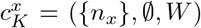, where *n*_*x*_ is a node in *N* indexed by *x* ∈ *ℕ*, and *K* ∈ ℕ is as defined in Lemma 7.1.
- **Inductive step**: The cluster 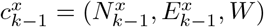 is defined as:
  – A node *n* is in 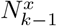 if there exists *m* ∈ ℕ and cluster indices *y*_1_, …, *y*_*m*_ ∈ *N* such that:
    1. *y*_1_ = *x*
    2. 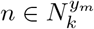
    3. For each *i* ∈ {1, …, *m* − 1}, there exists an edge *e* ∈ *T*_*k*−1_ with one endpoint in 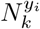 and the other in 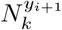
  – An edge *e* is in 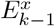 if both endpoints of *e* are in 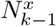 and *e* ∈ *H*_*k*−1_

Note that when clusters 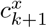 and 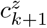 merge at step *k*, we have:

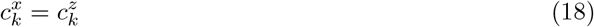

Furthermore, note that *m* = 1 is permitted, in which case the condition is satisfied by individual nodes with a path with no necessary edges. From this, we conclude 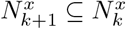.

The proof strategy now is to prove the clusters produced by the IC-TFCE are MCIS. As the MCIS from a subgraph *G*_*k*_ are unique and the TFCE computes the MCIS of subgraphs *G*_*k*_, if the IC-TFCE computes the MCIS of *G*_*k*_, they will be consequently equal from their uniqueness.

#### Lemma 7.5.

*The clusters* 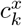 *constructed by the IC-TFCE are maximally connected induced subgraphs*

*Proof*. To prove the nodes and edges are maximal and the graph is connected, one can use reverse induction from *K* to 0. If those three requirements are satisfied, one has that the cluster is a MCIS by the definition of MCIS.

Let 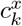 be a cluster generated by the IC-TFCE algorithm.

### Base case

From Lemma 7.1, one has that the set of edges is empty for the graph *G*_*K*_. Therefore, as the clusters are initially defined containing the nodes alone, the clusters in 𝒞_*K*_ are indeed maximum connected induced subgraphs.

### Induction Step

- **Nodes are Maximal**: Assume by contradiction that the number of nodes is not maximal: there exists a node 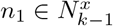 with an edge *e* = (*n*_1_, *n*_2_) ∈ *E*_*k*−1_ such that 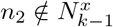. Since *e* ∈ *E*_*k*−1_ = *H*_*k*−1_ (Lemma 7.2) and *E*_*K*_ = ∅, there exists some *K* > *i* ≥ *k* − 1 such that *e* ∈ *T*_*i*_. At threshold *i*, let 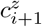 be the cluster containing *n*_1_ and let 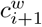 be the cluster containing *n*_2_. Since *e* = (*n*_1_, *n*_2_) ∈ *T*_*i*_, the edge *e* is a bridging edge between 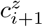 and 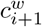 at step *i*. This implies 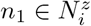 and 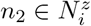 and they are both in the same cluster (indexed with *z*). By cluster construction, one has that 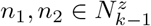 as 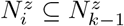. Since 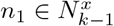, from the definition in Equation (18), one has that 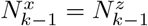 and that 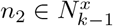 a contradiction to the assumption that 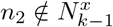. It follows that the number of nodes is maximal.
- **Edges Are Maximal**: Let us assume by contradiction that the number of edges is not maximal: there exists an edge *e* from 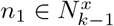 to 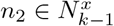 such that 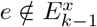. To avoid a contradiction with the algorithms definition, one must have *e* ∉ *H*_*k*−1_. However, from Lemma 7.2, *H*_*k*−1_ = *E*_*k*−1_, so if *e* ∉ *H*_*k*−1_, it is not in the edge *E*_*k*−1_ set of the subgraph *G*_*k*−1_. Since the cluster is from the subgraph *G*_*k*−1_ at iteration *k* − 1 within the set of edges *E*_*k*−1_, this is a contradiction and no such *e* is outside of this scope of edges, and consequently, the number of edges is maximal.
- **Graph is connected**: Let two nodes in 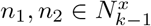. From the definition of the set of nodes for each cluster in iteration of the algorithm, for *n*_1_ to be in 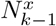, there exists a node 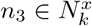 (can be *n*_1_) that is connected to *n*_1_ through a path of bridging edges from the previous iteration step. Again, repeating the same argument, from the definition of the algorithm, there exists a node 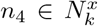, such that there is a path from *n*_2_ to *n*_4_. Since, one has that 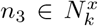 and 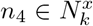, there exists a path from *n*_3_ to *n*_4_ as 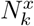 as is a set of nodes from the cluster 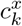 must be a connected graph from the induction assumption.

Therefore, the path:

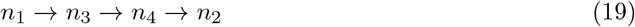

proves that the nodes *n*_1_ and *n*_2_ are connected by a path.

With this induction, one can conclude 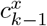 is a MCIS and the proof is concluded. □

From the results from Corollary 7.3, we establish that the edge allocation procedure from the IC-TFCE creates subgraphs equal to the TFCE. On Lemma 7.5, we establish that the clusters from IC-TFCE are indeed MCIS. Since its folklore that MCIS are unique for a given graph, it follows that IC-TFCE generates the same clusters as the traditional TFCE implementation.

### 7.1 Extension to Exact TFCE

It suffices to propose *T*_*k*_ such that

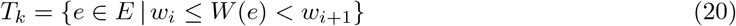

where (*w*_*i*_)_*i*∈ℕ_ is the ordered (monotonically increasing) sequence of weights for all images of *W*(*e*) with *e* ∈ *E*. The conclusions and reasoning from proofs from the IC-TFCE would follow.

## 8 Voxel Graph Transformation

In this section, we prove that the graph-based transformation yields clusters equivalent to the traditional voxel cluster definition used in neuroimaging.

### 8.1 Voxel Data and Traditional Clusters

Let 𝒱 be a finite set of voxels and let ~ denote spatial adjacency (voxels neighboring each other) on 𝒱, where we include self-adjacency (i.e., *v* ~ *v* for all *v* ∈ 𝒱). Let *T* : 𝒱 → R be a function mapping each voxel to its test statistic. We refer to the triple (𝒱, ~, *T*) as *voxel data*.

Given a threshold *h* ∈ ℝ, a voxel *v*_*x*_ ∈ 𝒱, and voxel data 𝒟 = (𝒱, ~, *T*), the thresholded cluster definition which is used by the voxel-based TFCE and cluster size inference in neuroscience:

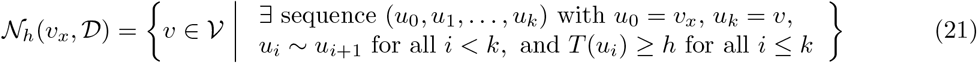

Intuitively this means that if you take any voxel in the cluster, you can recreate the cluster by using adjacent paths above the desired threshold.

### 8.2 Graph Transformation

We define the equivalent graph transformation Γ that maps voxel data to a weighted graph. Given voxel data 𝒟 = (𝒱, ~, *T*), define Γ : (𝒱, ~, *T*) → *G* = (*N, E, W*) where:

#### Nodes *N*

Each voxel becomes a node in the graph:

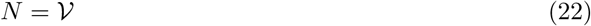

#### Edges *E*

Two nodes are connected by an edge if and only if their corresponding voxels are spatially adjacent. Recall that we include self-adjacency (*v* ~ *v* for all *v*), so each node also has a self-loop:

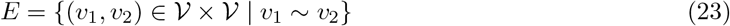

#### Weights *W*

The weight function maps an edge to the minimum test statistic of its two endpoints:

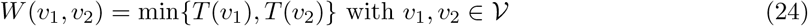

Note that the self-adjacency condition ensures that the edges from a node to itself (*v, v*) ∈ *E* result in *W*(*v, v*) = *T*(*v*) for each voxel.

### 8.3 Graph-Based Clusters

Given a threshold *h* ∈ ℝ, define the following subset of edges from *E*:

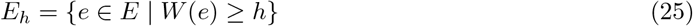

For a node *v*_*x*_ ∈ *N* and voxel data 𝒟, define the *graph cluster* 𝒮_*h*_(*v*_*x*_, Γ(𝒟)) as the set of nodes reachable from *v*_*x*_ via edges in *E*_*h*_:

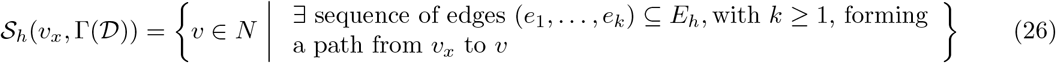

### 8.4 Equivalence of Cluster Definitions

#### Theorem 8.1.

*Let G* = Γ(𝒟) *be the equivalent graph transformation applied to voxel data* 𝒟 = (*V*, ~, *T*). *For any threshold h* ∈ ℝ *and any voxel v*_*x*_ ∈ 𝒱:

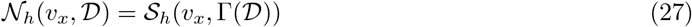

*Proof*. One can show that the sets are equal through mutual inclusion.

- **Let** *v* ∈ *S*_*h*_(*v*_*x*_, Γ(𝒟)): From the definition of 𝒮_*h*_(*v*_*x*_, Γ(𝒟)), there exists a path of edges in *E*_*h*_ from *v*_*x*_ to *v*, corresponding to a sequence of voxels (*u*_0_, *u*_1_, …, *u*_*k*_) with *u*_0_ = *v*_*x*_ and *u*_*k*_ = *v*. For each edge (*u*_*i*_, *u*_*i*+1_) in this path, one has *W*(*u*_*i*_, *u*_*i*+1_) ≥ *h*, which implies min{*T*(*u*_*i*_), *T*(*u*_*i*+1_)} ≥ *h*. Consequently, for all *i* ≤ *k*, it is *T*(*u*_*i*_) ≥ *h*. Since each (*u*_*i*_, *u*_*i*+1_) ∈ *E*_*h*_ ⊆ *E*, it follows from the definition of *E* that *u*_*i*_ ~ *u*_*i*+1_ for all *i < k*. Considering the following statements: it is *u*_0_ = *v*_*x*_, *u*_*k*_ = *v*, and *T*(*u*_*i*_) ≥ *h* for all *i* ≤ *k*, the sequence (*u*_0_, …, *u*_*k*_) satisfies all conditions in the definition of *N*_*h*_(*v*_*x*_, 𝒟), so *v* ∈ *N*_*h*_(*v*_*x*_, 𝒟).
- **Let** *v* ∈ *N*_*h*_(*v*_*x*_, 𝒟): From the definition of *N*_*h*_(*v*_*x*_, 𝒟), there exists a sequence of adjacent voxels (*u*_0_, *u*_1_, …, *u*_*k*_) with *u*_0_ = *v*_*x*_, *u*_*k*_ = *v*, and *T*(*u*_*i*_) ≥ *h* for all *i* ≤ *k*. Since *u*_*i*_ ~ *u*_*i*+1_, the edge (*u*_*i*_, *u*_*i*+1_) ∈ *E* for all *i* ≤ *k*. Since one has both *T*(*u*_*i*_) ≥ *h* and *T*(*u*_*i*+1_) ≥ *h* for each consecutive pair, one has:

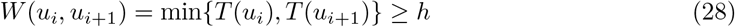

From (*u*_*i*_, *u*_*i*+1_) ∈ *E* and Equation (28), it follows from (*u*_*i*_, *u*_*i*+1_) ∈ *E*_*h*_ for all *i* ≤ *k*. It follows there exists a path in *E*_*h*_ from *v*_*x*_ to *v*, so *v* ∈ 𝒮_*h*_(*v*_*x*_, Γ(𝒟)).

One can conclude that the sets are indeed equal. □

